# NeuroSimo: an open-source software for closed-loop EEG- or EMG-guided TMS

**DOI:** 10.1101/2025.04.05.647342

**Authors:** Olli-Pekka Kahilakoski, Kyösti Alkio, Oula Siljamo, Kim Valén, Joonas Laurinoja, Lisa Haxel, Matilda Makkonen, Tuomas P. Mutanen, Timo Tommila, Roberto Guidotti, Giulia Pieramico, Risto J. Ilmoniemi, Timo Roine

**Affiliations:** Department of Neuroscience and Biomedical Engineering, Aalto University, Finland; A.I. Virtanen Institute for Molecular Sciences, University of Eastern Finland, Finland; Hertie Institute for Clinical Brain Research, University of Tübingen, Germany; Department of Neurology & Stroke, University of Tübingen, Germany; Machine Learning in Science, Excellence Cluster Machine Learning, University of Tübingen, Germany; Tübingen AI Center, University of Tübingen, Germany; Department of Neuroscience, Imaging and Clinical Sciences, Gabriele d’Annunzio University of Chieti and Pescara, Italy; Institute for Advanced Biomedical Technologies, University ‘G. D’Annunzio’ of Chieti-Pescara, Italy; Department of Engineering and Geology, University ‘G. D’Annunzio’ of Chieti-Pescara, Italy

**Keywords:** brain-state-dependent, brain stimulation, neurofeedback, neuromodulation, Python, real-time processing, ROS 2

## Abstract

**Objective:** Our goal was to create open-source software for closed-loop EEG–TMS that allows researchers to rapidly prototype and develop novel stimulation paradigms in a high-level programming language. This addresses the limitations of current solutions, which often rely on proprietary hardware and software, limiting their accessibility and customizability, or comprise ad-hoc pipelines tailored to specific use cases.

**Approach:** We developed NeuroSimo, a software platform that enables arbitrary EEG–TMS stimulation protocols written in Python, leveraging Python’s ecosystem of scientific, neuroimaging, and machine learning libraries. The core software is written in C++ with an embedded Python interpreter and employs the Robot Operating System (ROS 2) for inter-process communication. NeuroSimo runs on real-time-enabled Linux Ubuntu, using LabJack T4 for pulse triggering, and supports two EEG device models (Bittium NeurOne, BrainProducts actiCHamp) and TMS devices that deliver pulses via trigger signals. The software includes a graphical user interface for configuration and performance monitoring, and supports GPU processing for neural network computations.

**Main results:** In brain-state-dependent stimulation using the Phastimate algorithm, which targets TMS pulses to the trough of sensorimotor μ-rhythm, NeuroSimo achieved a median timing error of 0.2 ms (95% CI: 0.2–0.2 ms), a 99th-percentile of 0.6 ms (0.6–0.6 ms), and a maximum of 1.4 ms.

**Significance:** As an open-source platform combining Python’s flexibility with real-time closed-loop EEG–TMS, NeuroSimo enables researchers to develop and implement novel therapeutic approaches, marking a significant advance in personalized brain stimulation.

## Introduction

Transcranial magnetic stimulation (TMS) is a well-established technique for treating brain disorders and advancing our understanding of brain function. However, its effectiveness is limited by diminishing results over time and variability in individual responses [1–4]. The principle of spike-timing dependent plasticity [5,6] suggests that synchronizing stimulation with specific brain states can enhance long-term efficacy of TMS. Brain-state-dependent TMS, which aligns TMS pulses with biomarkers of ongoing brain activity derived from electroencephalography (EEG), has been shown to more effectively induce long-term potentiation (LTP)-like changes than traditional, unsynchronized methods [7,8]. This personalized, state-aware approach could mitigate current limitations of TMS, leading to more reliable and lasting therapeutic outcomes.

Closed-loop stimulation adapts in real time based on feedback from brain activity changes induced by stimulation, enabling dynamic and potentially more effective interventions than brain-state-dependent TMS [9]. Feedback is typically derived from EEG or electromyography (EMG). EEG directly measures brain activity from the scalp, while EMG indirectly reflects brain activity through muscle responses. Both techniques are well-established, non-invasive, and low-cost, promoting their use for real-time feedback in closed-loop systems.

Developing systems for closed-loop stimulation poses significant technical challenges because feedback must be processed within milliseconds to ensure timely adjustment of stimulation. Integrating realistic electric field models [10] or advanced prediction models, such as deep neural networks [11], further complicates these systems due to their high computational demands, often necessitating graphics processing unit (GPU) processing.

To ensure real-time operation, current systems sometimes execute stimulation algorithms outside the operating system (OS), as exemplified in [7]. However, this approach excludes GPU processing, which relies on OS support, and limits access to software libraries needed for modern machine learning algorithms. Commercial closed-loop systems, such as Sync2Brain’s bossdevice [12] or neuroCare’s LOOP-IT^1^, rely on specialized hardware that also typically lacks GPU compatibility. While well-suited for clinical use, the proprietary and restrictive nature of these systems may hinder research and experimentation unless substantial vendor support is available. The shift toward software-defined systems, as seen in the software-defined everything paradigm [13], offers a pathway to more flexible and accessible research tools.

An open-source alternative is The Brain Electrophysiological recording and STimulation (BEST) toolbox [14], a MATLAB-based library that supports real-time data processing and triggering of stimulation devices. However, achieving proper real-time performance with MATLAB often requires additional tools, such as Simulink Real-Time or RT-LAB, as MATLAB is not optimized for real-time execution [15]. Integrating advanced computational models that require GPU acceleration into these environments is difficult due to their limited support for modern scientific computing libraries. Another solution is Falcon [16], which is based on a C++ architecture for high performance. It allows users to define custom processing pipelines using a graph-based framework. However, adding new processing functionality necessitates expertise in C++ programming, which can make Falcon less accessible to researchers without extensive programming skills.

To address these limitations, we introduce NeuroSimo—an open-source platform designed for flexibility and real-time performance. By leveraging real-time-enabled Linux and running on commercial-off-the-shelf hardware, NeuroSimo offers cost-effectiveness, high performance, and rapid availability [17]. It supports custom EEG-preprocessing and stimulation algorithms written in Python, a language widely used in scientific computing [18], and includes a web interface that enables rapid prototyping and iterative development. These features make NeuroSimo a versatile and powerful platform for brain stimulation research.

## Design

NeuroSimo leverages a microservice architecture, a widely used approach for building flexible and scalable software [19]. This design divides the system into independently executed services that communicate through well-defined interfaces, allowing time-critical processes to run with higher priority, optimizing resource use [20]. In a closed-loop EEG–TMS system, stages such as data acquisition, preprocessing, and stimulation control naturally form distinct microservices, providing a straightforward mapping from the problem space to the technical implementation, effectively addressing the problem of defining microservice boundaries [21]. Microservice approach also offers outside observability at checkpoints between microservices [22], such as after data preprocessing. Moreover, performance bottlenecks can be more readily identified due to each stage forming an independent unit, which can be measured and optimized separately [23].

NeuroSimo uses the Robot Operating System (ROS 2) as its communication infrastructure [24], selected for its suitability for real-time applications [25,26] and support for versatile communication patterns. While originally designed for robotics, these features and ROS 2’s ability to interface with diverse hardware make it an excellent choice for building software beyond its initial scope. NeuroSimo’s core is implemented in C++ to ensure the determinism and high performance required for real-time applications [27], with Python integration achieved through pybind11^2^, enabling custom Python algorithms to run seamlessly within the C++ process.

In ROS 2, processing is divided into units called nodes, each corresponding to a microservice in NeuroSimo’s design. These nodes use ROS 2’s publisher-subscriber communication model for inter-service messaging and employ higher-level communication patterns, such as ROS 2’s services and actions. In addition, ROS 2’s ability to retain the latest published messages^3^, effectively forming a small-scale database, is leveraged to persist state across microservice restarts.

The microservices are run in Docker containers [28], with one microservice per container, ensuring isolation and avoiding Python dependency conflicts [29], which is particularly beneficial for complex machine learning and EEG processing libraries. Each service maintains its own log, enabling efficient system state tracking and troubleshooting [30]. Docker Compose manages the containers [31], while the built-in Linux tool systemd ensures they start automatically at system boot [32].

NeuroSimo runs on a real-time-enabled Linux Ubuntu operating system to minimize delays in time-critical processes [33], employing Linux’s real-time processing capabilities. Time-critical services are assigned real-time priorities to prevent delays caused by lower-priority tasks, while best practices such as memory locking further reduce timing variability [34]. A web user interface, built with ReactJS [35], communicates with the system using rosbridge_suite and roslibjs^4^.

These design choices—microservice architecture, ROS 2 communication, Docker containerization, and integration of real-time capabilities—equip NeuroSimo with the flexibility, scalability, and performance required for rapid response to varying brain states.

### Overview of the Two-Module Design

NeuroSimo adopts a two-module design that decouples the computational decision-making process from the timing mechanism (Fig. 1). The Decision-Making Module processes sliding windows of EEG data to determine whether and when stimulation should occur, either for each new EEG sample or at specified intervals. The Timing Module ensures that pulses are triggered precisely at the specified times, independent of variability in decision-making computation. This separation maintains timing precision, even unexpected delays occur in the decision-making process. The two modules function similarly to a planner and an actuator: the planner identifies optimal stimulation times, while the actuator executes the plan with high precision.

**Figure 1:**
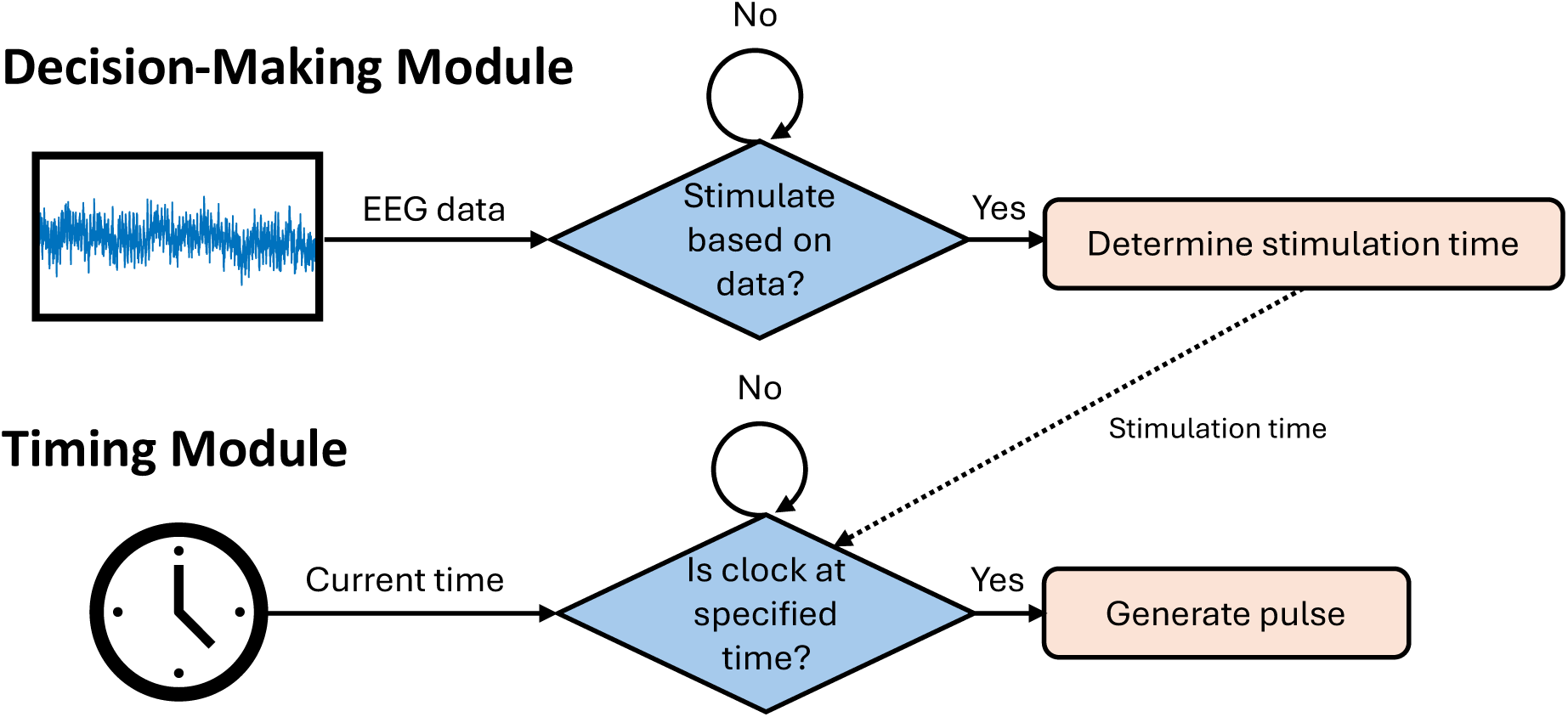
Flowchart illustrating the two-module design. The Decision-Making Module analyzes sliding windows of EEG data to determine whether and when to stimulate. If stimulation is required, the desired time is sent to the Timing Module, which ensures the stimulation pulse is generated precisely at the specified time, unaffected by variability in the execution time of the Decision-Making Module.

In contrast to a straightforward single-module design that combines decision-making and timing, NeuroSimo divides these responsibilities, ensuring a strong separation of concerns [36] and safeguarding timing accuracy from computational fluctuations in decision-making.

This enhances system reliability and supports potential future extensions, such as refining stimulation decisions post hoc based on additional information.

To enable the Timing Module to generate pulses based on timing information from the Decision-Making Module, both must share a common notion of time. Ideally, this could be achieved if the EEG, TMS, and control device were synchronized to a common clock, but current EEG and TMS devices typically lack support for clock synchronization protocols such as Precision Time Protocol [37]. To address this, NeuroSimo uses the EEG device clock as the reference, with the Timing Module estimating the current time based on the latest EEG sample. This approach keeps both modules synchronized and operating within the same time domain. Between samples, the Timing Module extrapolates time using its internal clock [38], ensuring precise timing even at low sampling rates.

Despite sharing a time reference, inherent delays—or latencies—in the system must be compensated for to achieve precise stimulation timing:

1. Timing latency (Δt_timing_) arises from the time required for the EEG sample to reach the Timing Module and for the pulse command to be delivered to the TMS device. If stimulation is planned at time *t*_0_, triggering a pulse when the EEG sample timestamped *t*_0_ is received would result in a delay. To compensate for this, the pulse is generated when the EEG sample corresponds to *t*_0_ − Δt_timing_.
2. Decision-Making Latency (Δt_decision_) refers to the delay between the Decision-Making Module receiving an EEG sample and the corresponding stimulation decision reaching the Timing Module. It also determines the earliest feasible stimulation time; for a stimulation decision based on an EEG sample at time *t*_0_, the earliest stimulation time is *t*_0_ + Δt_timing_ + Δt_decision_. To enable feasible stimulation decisions by the Decision-Making Module, both latencies are continuously monitored, and the module is updated with the estimated earliest stimulation time for each new sliding window.

Timing latency is measured with trigger signals, commonly used for rapid device-to-device communication. These signals can be generated from a PC using a peripheral triggering device. The Timing Module generates triggers at specific reference clock times, timestamps them, and calculates the time differences. If the triggering device has dual-output capability, one output can be used to timestamp events with the EEG device clock while the other triggers TMS pulses (Fig. 2). This setup decouples pulse generation from latency measurement, allowing continuous and automatic latency tracking during operation, provided that the TMS device delivers pulses promptly in response to triggers. In contrast, decision-making latency is measured internally by the PC, as it does not involve communication with external devices.

**Figure 2:**
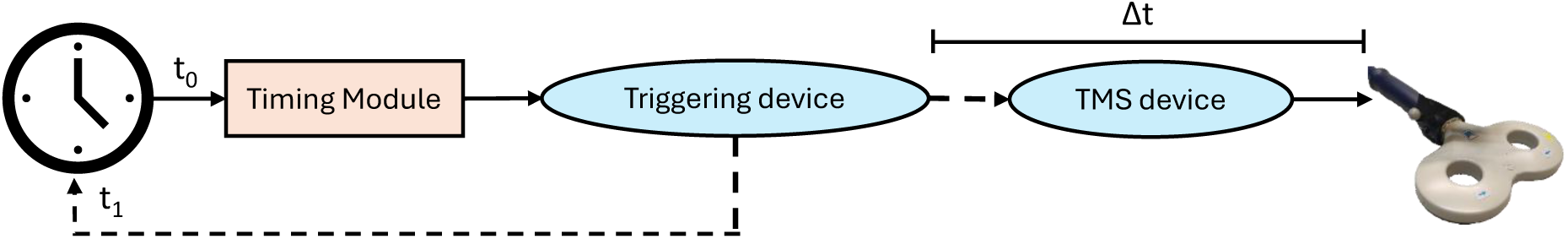
Measuring the latency of the Timing Module using triggers (dashed lines). The triggering device outputs two triggers: one is timestamped with the reference clock and the other triggers TMS pulses. Latency is the time difference between initiating the trigger (*t*_0_) and receiving it back (*t*_1_). It approximates the end-to-end latency between the timing module and the pulse delivery accurately provided that the trigger-to-pulse delay *Δt* is negligible.

Timing latency is influenced by specific PC hardware, including its interfaces with the EEG and triggering devices. Factors such as network integrity further impact latency [39]. To ensure an up-to-date estimate, NeuroSimo periodically re-measures timing latency (at 10 Hz) and automatically compensates for it using the latest measurement, exemplifying dynamic latency management (cf. [26]). By avoiding low-pass filtering of the measurements, the system can adapt rapidly to changes, keeping the compensation scheme responsive, straightforward, and avoiding the use of additional state information. Latency is continuously monitored, ensuring it remains within bounds and verifying the operation of the pipeline and correct PC configuration.

Precise pulse timing is ensured by monitoring timing errors via feedback from the TMS device (Fig. 3). The TMS device is configured to generate a trigger with each pulse, allowing the EEG device to timestamp these triggers directly using its reference clock. This real-time feedback enables NeuroSimo to assess actual pulse timing and verify its alignment with the EEG signal.

**Figure 3:**
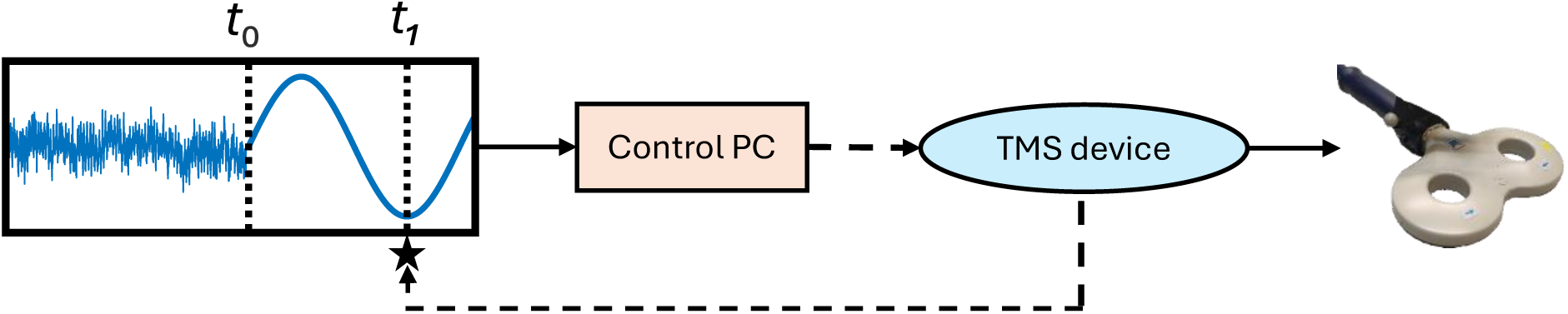
Real-time monitoring of pulse timing errors. The control PC analyzes a sliding window that contains EEG data up to the current timepoint t0 (left side of the window) and extends into the future with predicted phase values (right side). The TMS device delivers a pulse targeting at t1 and generates a simultaneous trigger (dashed line) that is sent back to the EEG device. This return trigger is timestamped with the reference clock (depicted by an asterisk) and forwarded to the control PC, enabling it to compute the timing error as the difference between the expected and actual pulse times.

### Overview of the Software Architecture

The software architecture of NeuroSimo is organized around two parallel data-processing pipelines that reflect its two-module design: the Timing Pipeline and the Decision-Making Pipeline (Fig. 4). Both pipelines use EEG data as input but diverge in functionality: the Timing Pipeline controls TMS pulse triggering at specified times, while the Decision-Making Pipeline processes EEG data to determine whether and when to stimulate, passing this information to the Timing Pipeline.

**Figure 4:**
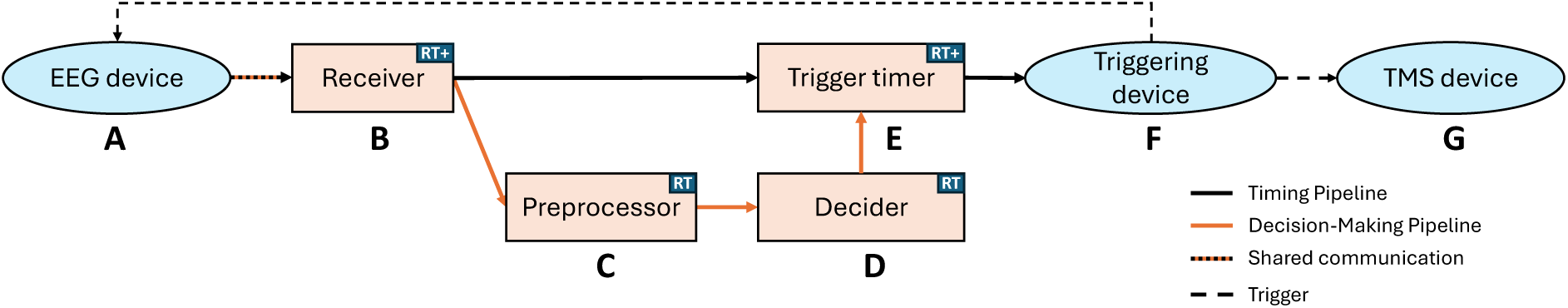
Architecture of the core pipelines, depicting devices (ellipses), microservices (rectangles), and their communication (arrows). Black arrows represent the Timing Pipeline, orange arrows represent the Decision-Making Pipeline, and black–orange arrows indicate shared communication between the pipelines. Dashed arrows depict triggers. “RT” denotes real-time priority, while “RT+” indicates an elevated real-time priority for prioritizing the timing pipeline.

The core architecture comprises several devices and microservices, each with a specific role:

1. EEG device (A): Acquires EEG signals and transmits them to the computer running NeuroSimo.
2. Receiver (B): Reads incoming EEG data and publishes it in a unified format for downstream services.
3. Preprocessor (C): Processes the EEG data using a Python script and publishes the preprocessed data.
4. Decider (D): Analyzes preprocessed data with a Python script to determine stimulation decisions and publishes them.
5. Trigger timer (E): Uses a triggering device to generate trigger signals at specified times.
6. Triggering device (F): Generates trigger signals for pulse delivery and latency measurement.
7. TMS device (G): Delivers stimulation pulses in response to trigger signals.

To support brain–computer interfaces or psychophysiological experiments, the system additionally includes a service called the Presenter. This module subscribes to stimulation decisions and delivers visual or auditory stimuli in response, enabling interaction with the subject based on ongoing brain activity patterns.

### Python and C++ Integration

NeuroSimo combines C++ for performance-critical tasks and Python for algorithmic flexibility, balancing performance with ease of development [40,41]. C++ handles computationally intensive tasks, such as maintaining sliding windows of past EEG samples. These sliding windows are passed to Python scripts for further processing, enabling the use of algorithms leveraging popular libraries such as NumPy [42] and SciPy [43], which rely on highly optimized C and Fortran code, or CuPy [44] and TensorFlow [45] for GPU processing (Fig. 5).

**Figure 5:**
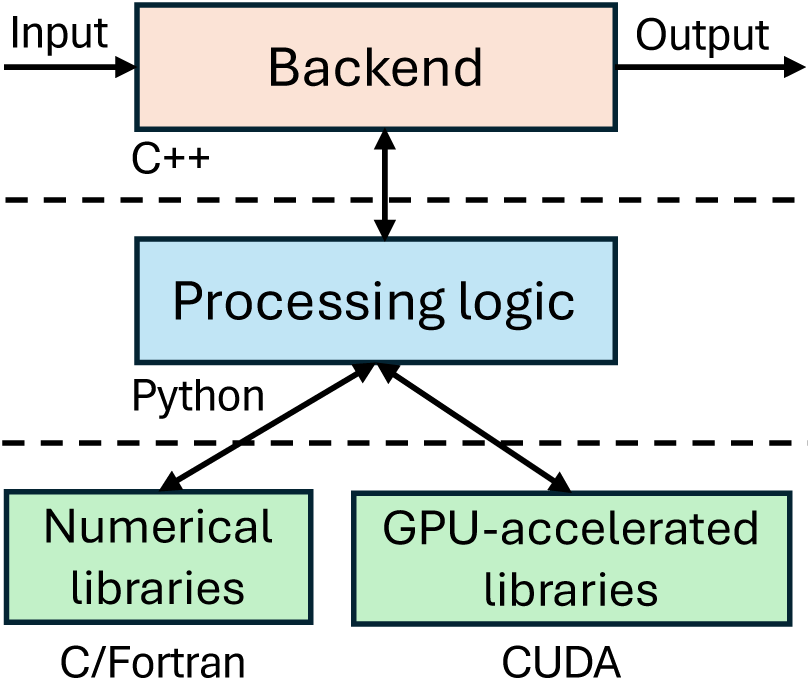
Interaction between C++ (top), Python (middle), and numerical and GPU-accelerated libraries (bottom). The C++ backend provides the infrastructure for tasks such as reading the EEG data and communicating with other modules. Processing logic is defined in Python, which can employ libraries such as SciPy and TensorFlow. These libraries utilize optimized C and Fortran routines or GPU acceleration for efficient numerical operations.

The Preprocessor and Decider modules communicate with their respective Python scripts regularly during system operation (Fig. 6). At initialization, the scripts receive information about the data stream, including the sampling rate and the number of EEG and EMG channels. They are also queried for their configurations, such as the desired length of the sliding window.

**Figure 6:**
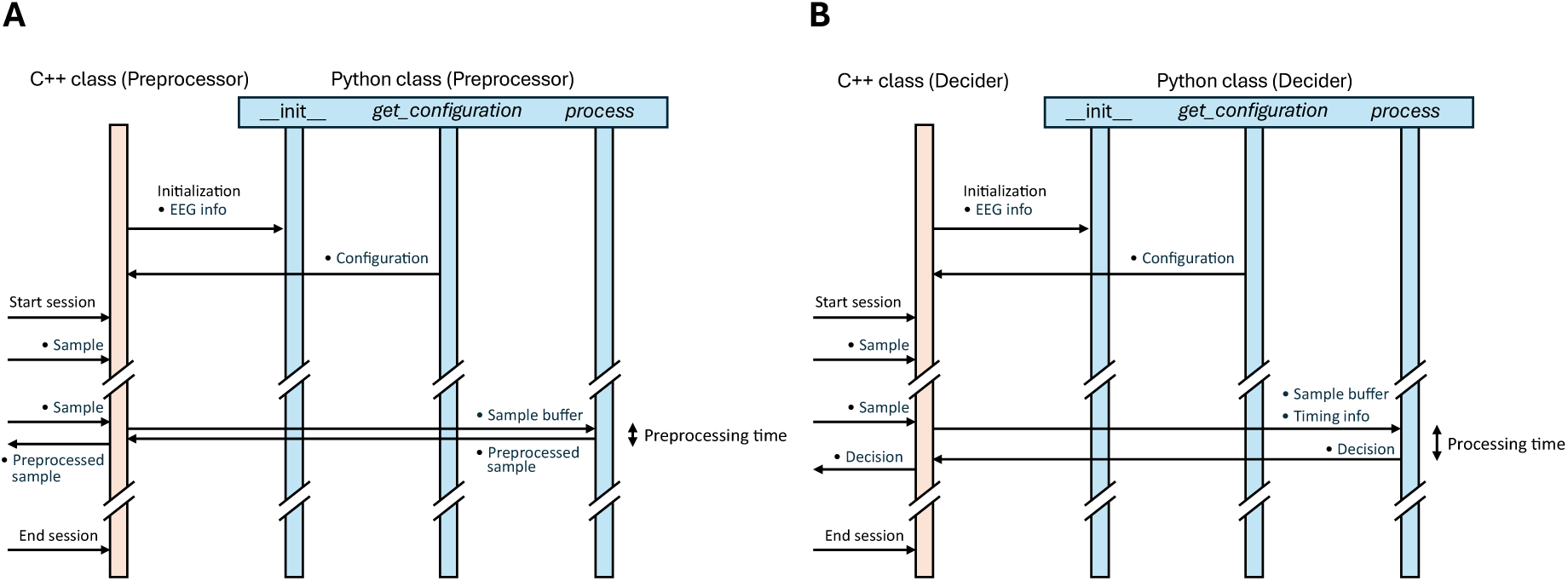
Sequence diagrams illustrating the communication between the C++ backend and the Python modules for the Preprocessor and Decider services (A and B). Bullet points indicate variables such as EEG samples and sliding windows. Both modules include methods for initialization, configuration queries, and sample processing. The Preprocessor Python script outputs a preprocessed sample for each received sliding window (A), while the Decider determines whether and when to stimulate based on the sliding window and timing information containing the estimated earliest stimulation time (B). Both backends only start passing sliding windows to the Python modules once these windows are full of samples, preventing the need to handle partial windows.

During streaming, sliding windows are passed to the process methods of the Preprocessor and Decider scripts. The Preprocessor outputs a single preprocessed sample for each sliding window, ensuring a continuous and steady stream of EEG data. Samples can be tagged as invalid, such as when excessive noise is detected, preventing their use in decision-making.

The Decider generates stimulation decisions at either specific intervals or predefined timepoints, depending on experimental requirements.

To optimize performance, sliding windows are implemented using circular buffers with a constant-time append operation [46]. Before passing the data to Python, the circular buffer is unwound into a linear buffer, creating a continuous data structure that enables efficient matrix operations [47]. By passing the buffer to Python by reference, the system minimizes memory copying, further improving performance [48]. Since the Python interpreter runs within the same C++ process, it automatically inherits the real-time priority assigned to the process, ensuring consistent responsiveness and seamless integration. As Python runs within the same C++ process, it inherits the process’s real-time priority, ensuring responsive performance.

Configurations, input arguments, and output formats for the Preprocessor and Decider scripts are detailed in the examples provided with the software, which also serve as templates for custom scripts.

### System setup

The system setup comprises an EEG device, control computer, triggering device, and TMS device (Fig. 7).

**Figure 7:**
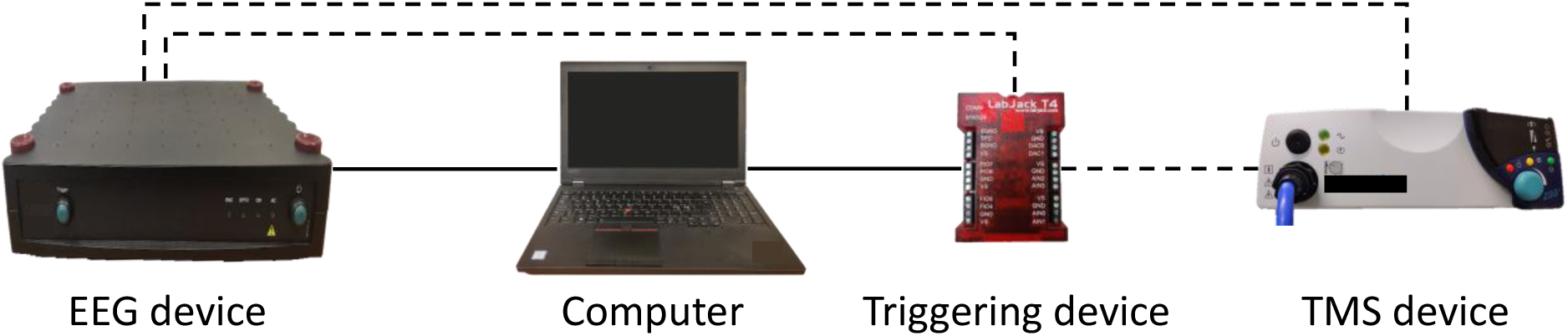
Device setup showing connections (lines) and triggers (dashed lines). The triggering device (LabJack T4) connects to both the TMS device for pulse delivery and the EEG device’s trigger-in port for latency measurement. A trigger line from the TMS device to the EEG device enables automatic measurement of timing errors, with the TMS device configured to generate a simultaneous trigger with each pulse.

NeuroSimo supports two EEG devices: Bittium NeurOne (Bittium Biosignals Ltd, Finland) and Brain Products actiCHamp (Brain Products Gmbh, Germany). Both transmit measurement data to the computer via the User Datagram Protocol (UDP), which is well-suited for real-time data transfer due to its low latency [49]. However, as UDP does not support retransmission of lost packets, the software must handle potential dropped samples.

The LabJack T4 USB peripheral serves as the triggering device, with a LJTick-DigitalOut5V module providing Transistor-Transistor Logic (TTL)-level compatibility. The module’s pull-down resistor prevents prevents accidental TMS triggering during system start-up or potential malfunctions. Triggers are generated as 1-millisecond pulses, which are reliably detected by both TMS and EEG devices.

NeuroSimo is compatible with any TMS device that supports pulse delivery in response to trigger signals. NeuroSimo also supports the multilocus TMS device developed at Aalto University [50,51], which achieves sub-millisecond precision in pulse timing by delegating timed pulse delivery to the TMS device itself, as detailed in [52].

### Web User Interface

NeuroSimo provides a web-based user interface for easy configuration, control, and monitoring (Fig. 8).

**Figure 8:**
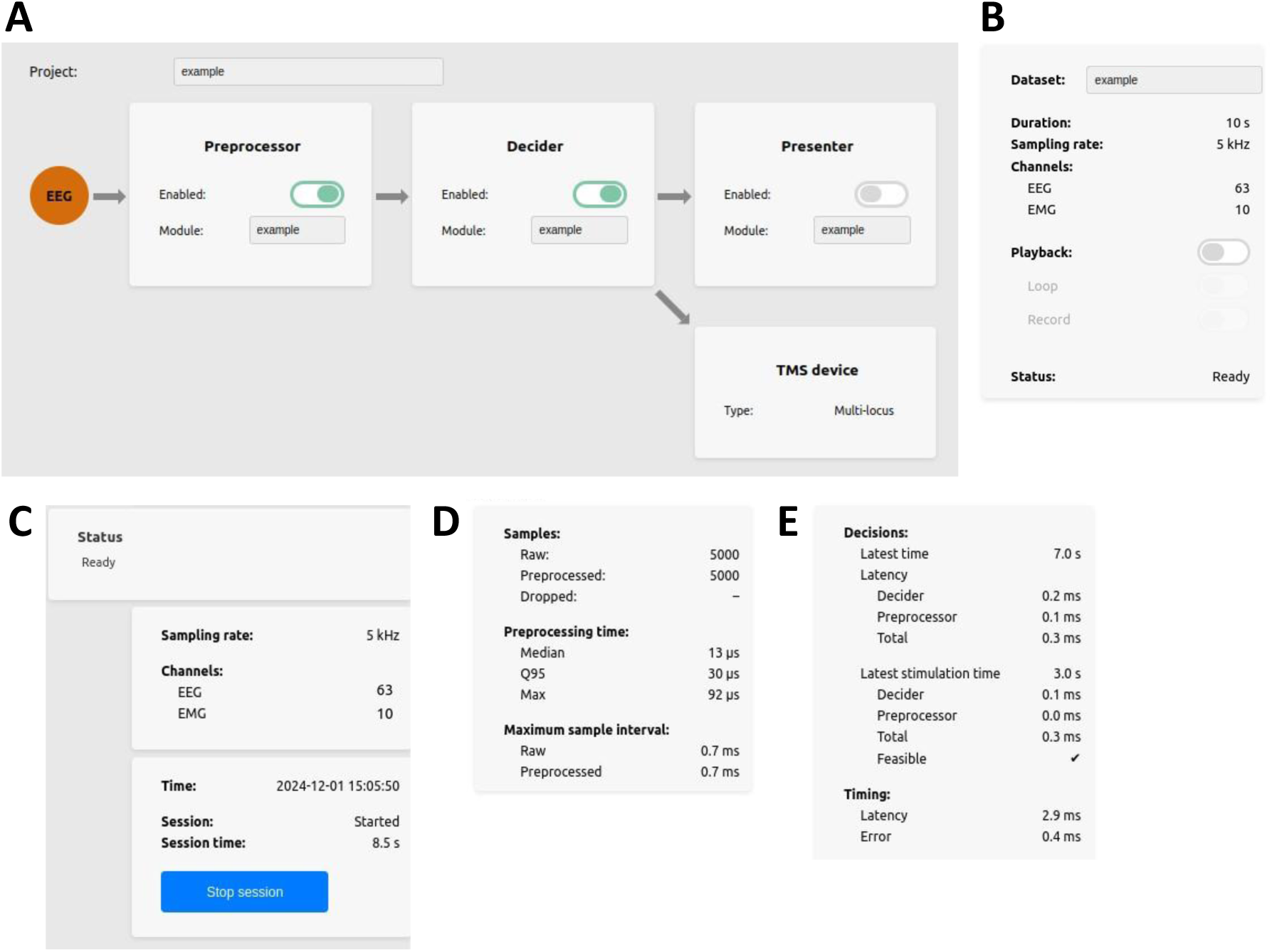
The web user interface: (A) Selection of the project and Python scripts. (B) Status bar, including EEG stream information and session controls (start and stop). (C) EEG simulator for streaming offline EEG data from pre-recorded datasets. (D) Statistics for system monitoring: sample counts, preprocessing times, and sample intervals, aggregated per second. (E) Latency profiles and feasibility for each stimulation decision, timing latency (updated periodically), and timing error (updated with each pulse).

The Python scripts for the Preprocessor, Decider, and optional Presenter modules can be selected via dropdown menus (Fig. 8A), and each module can be independently enabled or disabled. For example, disabling the Preprocessor routes raw EEG data directly to the Decider, allowing testing different configurations and simplifying the pipeline.

NeuroSimo features an EEG simulator (Fig. 8B) that plays back pre-recorded EEG data in CSV format for testing and simulation. Each dataset is accompanied by a JSON file containing metadata, such as the dataset name. The system also supports seamless repetition of datasets, allowing even short EEG recordings to be used for prototyping and algorithm development.

The user interface displays real-time statistics, such as sample count and preprocessing time per sample, aggregated every second (Fig. 8D). Stimulation decisions are profiled for their latency and timing feasibility. The timing latency of the system is updated periodically, and the timing error of each successful pulse is presented, enabling continuous monitoring of the system performance and quick detection of issues (Fig. 8E).

The interface supports rapid prototyping and live debugging of preprocessing and decision-making algorithms, with the scripts automatically reloading after modifications without requiring session restarts. New Python scripts are automatically detected and made available in the drop-down menu for easy integration.

### Error Handling and Safeguards

NeuroSimo includes safeguards and error-handling mechanisms to ensure safety and reliability, aligning with the layered safety approach [53] essential in safety-critical applications. For example, to adhere to TMS safety guidelines [54], a configurable minimum intertrial interval is enforced (default: 2 seconds). If stimulation is initiated more frequently, additional pulses are prevented, and a warning is issued. This safeguard does not affect pulse pairs, which, if supported by the TMS device, are typically generated internally in response to a single trigger.

Another safeguard addresses pipeline performance: if the decision-making pipeline cannot keep up with the incoming data stream, stimulation is automatically disabled with an error message displayed. Users can then optimize the pipeline or reduce the decision-making script’s calling frequency.

Stimulation is automatically disabled if the timing latency exceeds a configurable threshold (default: 5 ms). If stimulation is requested at a time that cannot meet these latency constraints, the pulse is prevented and a warning is issued.

Dropped EEG samples are continuously monitored, with the cumulative count displayed in the user interface. If the drop rate exceeds a configurable limit (default: 4 samples per second), stimulation is automatically disabled. This safeguard identifies issues such as compromised network connections, ensuring reliable input data for decision-making.

### Experimental verification

NeuroSimo’s performance was evaluated on a desktop workstation (Intel Core i7-13700K, GeForce RTX 3080) using two stimulation algorithms: “Trigger Once in a Second” (TOIS) and Phastimate [55], a phase-prediction method for targeting specific phases of brain oscillations. TOIS served as a baseline for assessing fundamental system characteristics such as timing latencies and errors, while Phastimate tested performance under realistic conditions.

In the Phastimate experiment, we recorded 45 minutes of resting-state EEG from a healthy, right-handed 34-year-old male volunteer in three batches. The human experiment was approved by the Coordinating Ethics Committee of the Hospital District of Helsinki and Uusimaa, adhered to the Declaration of Helsinki, and the participant provided written informed consent. In the TOIS experiment, disconnected EEG electrodes were used, and no human participants were involved. Similarly, 45 minutes of data were recorded.

EEG recordings were conducted at a 5 kHz sampling rate using a Bittium NeurOne system (Bittium Biosignals Ltd, Finland) and a TMS-compatible 62–channel cap (EasyCap BC-TMS-64-X20-UCMW, EasyCap, Herrsching, Germany) with Ag/AgCl sintered C-electrodes, positioned according to the International 10–20 system. The reference electrode was placed on the right mastoid, and the ground electrode on the right cheekbone, with electrode impedances below 5 kΩ. Raw data acquired at 80 kHz was low-pass filtered at 1250 Hz and downsampled to 5 kHz by the NeurOne system.

To maintain a realistic experimental setup, TMS pulses were delivered in response to triggers using a MagStim Rapid^2^ system (Magstim Company Ltd, Wales). However, the pulses were not delivered to the human subject to avoid TMS-induced artifacts that interfere with the post-hoc analysis of the actual triggered phase. The MagStim system was configured to generate with each pulse a simultaneous output trigger, connected to the NeurOne system to record precise pulse timing for post-hoc analysis of triggering accuracy and automatic measurement of timing errors.

In Phastimate, 1-second sliding windows at 1-second intervals were used to predict the future phase (36 ms ahead) of the μ-rhythm from the C3-Hjorth channel using an autoregressive model applied to band-pass filtered data (center frequency: 10 Hz), following [55]. The filtered signal was Hilbert-transformed to obtain an analytic representation, from which instantaneous phases were extracted. A trigger was generated if any predicted future phase fell within 0.05 radians of the trough, targeted at the corresponding future time point. If no suitable future phases were found, the trigger was skipped. In TOIS, stimulation was always timed 5 ms into the future.

Latencies and timing errors were measured using NeuroSimo’s functionalities and analyzed in R (version 4.4.1). The Phastimate algorithm was implemented following [55]. Confidence intervals for timing measurements were computed using bootstrap with 10,000 repetitions.

Post-hoc analysis in MATLAB (R2021b) provided an independent means to verify NeuroSimo’s timing accuracy and the Phastimate implementation. The instantaneous phases were extracted from the measured EEG signal in a manner similar to the online analysis, and these phases were used to investigate the alignment of the actual troughs with trigger times recorded by the NeurOne system.

## Results

Fig. 9 presents pooled timing latencies (A), decision-making latencies (B) for both algorithms, total latencies (C) for both algorithms, and pooled timing errors (D). Unless otherwise noted, medians and percentiles had 95% confidence intervals within 0.1 ms of the reported values. Timing errors were comparable between algorithms (Wilcoxon rank sum test, p = 0.99), with a median of 0.2 ms and a 99th percentile of 0.6 ms. Only 0.3% (0.1–0.5%) of errors exceeded one millisecond, with a maximum error of 1.4 ms across both 45-minute experiments.

**Figure 9:**
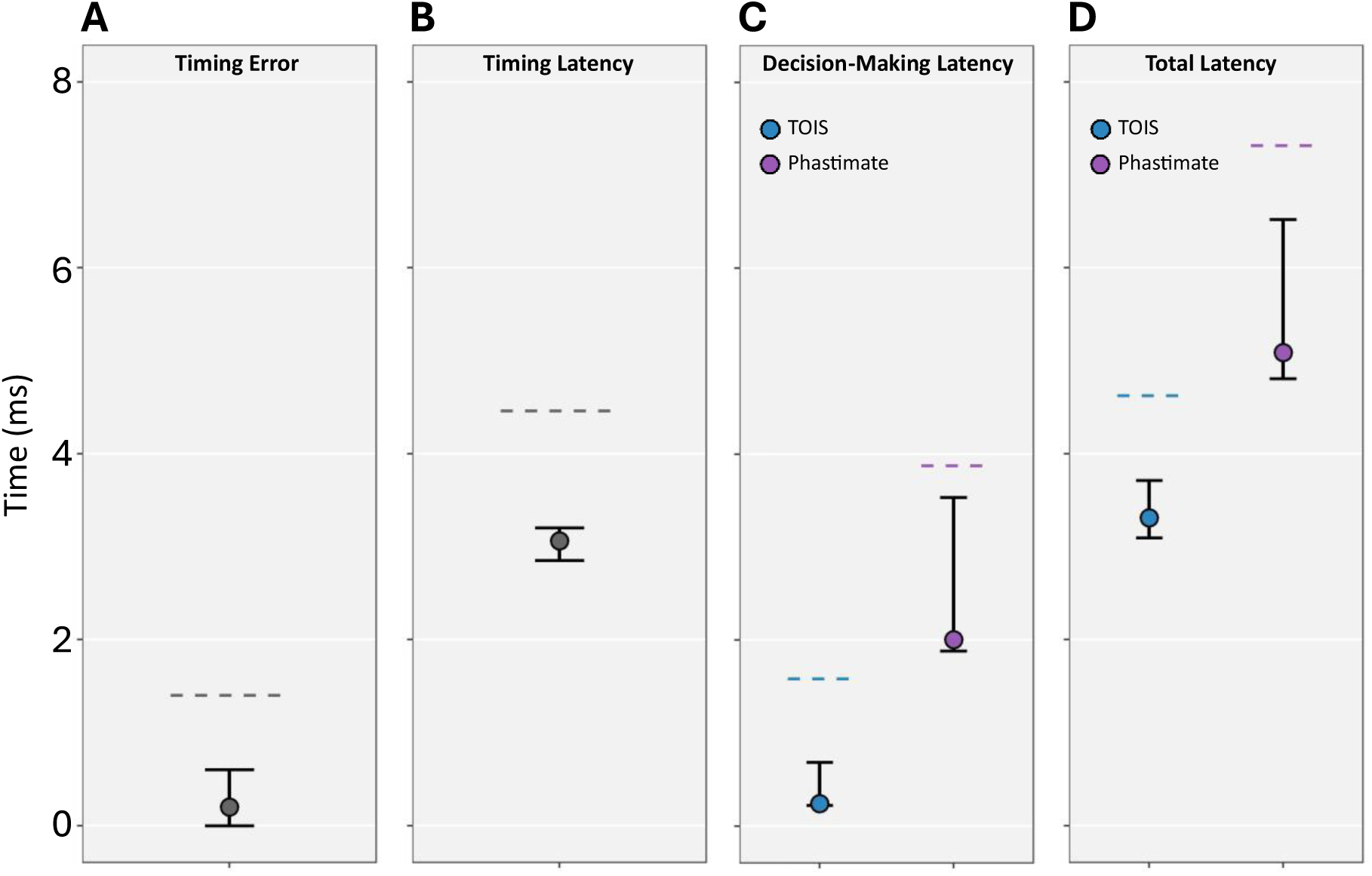
(A) Timing Error: pooled results for both algorithms. (B) Timing Latency: pooled results, as in (A). (C) Decision-Making Latency: TOIS (blue) and Phastimate (purple). (D) Total Latency: TOIS (blue) and Phastimate (purple). In each plot, the median is shown as a circle, whiskers represent the 1st–99th percentiles, and the horizontal line marks the maximum across the experiment.

Median timing latency was 3.1 ms, with a 99th percentile of 3.2 ms, showing consistent real-time performance. Decision-making latency was significantly longer for Phastimate (median: 2.0 ms) than TOIS (median: 0.2 ms), reflecting the computational demands of phase prediction. The total latency had a median of 5.1 ms and a 99th percentile of 6.5 ms (6.3–6.7 ms) for Phastimate, and a median of 3.3 ms and a 99th percentile of 3.7 ms (3.5–4.0 ms) for TOIS.

During the 45-minute Phastimate experiment, 2705 EEG data windows were processed for potential stimulation. A total of 673 pulses (25% of the windows) were triggered based on finding a suitable phase within the prediction window. Post-hoc analysis of the μ-rhythm trough targeting showed a mean phase error of 10° (4–17°), verifying the accuracy of phase prediction.

These findings demonstrate that NeuroSimo achieves timing errors in low millisecond range under realistic conditions.

## Discussion

NeuroSimo is a free and open-source software designed for closed-loop EEG–TMS, enabling experiments with commercial-off-the-shelf hardware and standard TMS devices. It allows researchers to define the logical flow of signal processing and decision-making in Python, providing a flexible and accessible programming interface.

The planner–actuator architecture, reflected in the two-module design, ensures robust stimulation timing and enables features such as planning stimulation into the future. By decoupling decision making from pulse timing, this architecture simplifies the conceptual framework for timing and ensures that timing precision is unaffected by delays in the decision-making process. In addition, it supports novel features such as automatic latency compensation, enabling researchers to focus on stimulation algorithms without the need to manually manage latency.

NeuroSimo combines real-time Linux, C++, and Python to offer the high performance of C++ alongside the flexibility of Python scripting. Recent advancements, such as the integration of real-time capabilities into the mainline Linux kernel^5^ and the introduction of a just-in-time (JIT) compiler in Python 3.13.0, further enhance the suitability of these technologies for real-time closed-loop TMS.

Automatic latency compensation in NeuroSimo enhances adaptability by enabling real-time adjustments to changing conditions. The system includes monitoring and safeguards to handle latency limits, ensuring reliable operation despite the complexity of adaptive timing.

The system supports combining a preprocessing script with a decision-making algorithm. Future extensions could enable chaining multiple preprocessing steps, inspired by modular audio signal processing, with configurable parameters such as filter cut-off frequencies and operations like downsampling and upsampling. Additional developments include expanding support for psychophysiological experiments and brain–computer interfaces, building on the foundation provided by the Presenter module.

While not yet implemented, the two-module architecture supports conditional or dynamically adjustable stimulation, such as the ability to abort or re-schedule pulses. Pulse timing could incorporate real-time adjustments to parameters such as waveform, intensity, and stimulation location, either via robot-guided or multichannel TMS. Implementing these features would require replacing hardware triggers with serial digital messages for stimulation control [52], likely in collaboration with TMS device manufacturers.

Future closed-loop EEG–TMS systems should aim for synchronized device clocks, with the TMS device handling precise pulse timing. This approach [52] would enable even better timing precision and robustness than what is achievable with the current method of control computer managing timing.

NeuroSimo is available via GitHub at https://github.com/neurosimo/neurosimo, with installation instructions and examples. We welcome contributions to expand documentation, improve functionality, and develop new features.

## Conclusion

NeuroSimo provides an open-source platform for closed-loop EEG–TMS experiments, combining reliable and precise timing with the accessibility of Python-based customization. The software enables researchers to implement and test new stimulation algorithms while maintaining precise temporal control of pulse delivery. It provides a foundation for development of new brain stimulation protocols.

## Acknowledgements

This work has been supported by the European Research Council (ERC Synergy) under the European Union’s Horizon 2020 research and innovation programme (ConnectToBrain; grant agreement No 810377), Research Council of Finland (grant agreement No 353798), US Global Program Pilot funding provided by Aalto University, Emil Aaltonen Foundation, the Finnish Cultural Foundation, and the Else Kröner Medical Scientists Kolleg Clinbrain: Artificial Intelligence for Clinical Brain Research. We thank Tuomas J. Lukka, Tomi-Mikael Kahilakoski, and Veli Peltola for useful discussions. OpenAI’s GPT-4 and Anthropic’s Claude 3.5 Sonnet were used as tools for revising the manuscript. All final content has been reviewed and approved by the authors.

## Author Contributions

Olli-Pekka Kahilakoski: Conceptualization, Methodology, Software, Validation, Formal Analysis, Writing – original draft, reviewing & editing. Kyösti Alkio: Software, Writing – reviewing & editing. Oula Siljamo: Software, Writing – reviewing & editing. Kim Valén: Software, Writing – reviewing & editing. Joonas Laurinoja: Software, Validation, Formal Analysis, Writing – reviewing & editing. Lisa Haxel: Software, Methodology, Writing – reviewing & editing. Matilda Makkonen: Methodology, Writing – reviewing & editing. Tuomas P. Mutanen: Methodology, Writing – reviewing & editing. Timo Tommila: Methodology, Writing – reviewing & editing. Roberto Guidotti: Software, Methodology, Writing – reviewing & editing. Giulia Pieramico: Methodology, Writing – reviewing & editing. Risto J. Ilmoniemi: Project administration, Supervision, Funding acquisition, Writing – reviewing & editing. Timo Roine: Conceptualization, Project administration, Supervision, Writing – reviewing & editing.

## Data Availability Statement

The NeuroSimo software is available at https://github.com/neurosimo/neurosimo. The timing measurements and analysis scripts used in the experimental validation are available on Zenodo at https://doi.org/10.5281/zenodo.14398633. The raw EEG recordings required for post-hoc analysis are not shared due to restrictions on sharing sensitive EEG data.

## Footnotes

1 Documentation for neuroConn’s closed-loop solution: https://info.neurocaregroup.com/hubfs/neuroCare_May_2021/pdf/neuroConn-LOOP-IT.pdf

2 pybind11: https://github.com/pybind/pybind11

3 ROS 2 Quality of Service: https://docs.ros.org/en/humble/Concepts/Intermediate/About-Quality-of-Service-Settings.html

4 rosbridge_suite software library: https://github.com/RobotWebTools/rosbridge_suite

5 PR MPT_RT was integrated into mainline Linux kernel in 2024.

## References

[1] Ridding MC, Ziemann U. Determinants of the induction of cortical plasticity by non-invasive brain stimulation in healthy subjects. Journal of Physiology 2010;588:2291–304. 10.1113/jphysiol.2010.190314.

[2] Lefaucheur JP, Aleman A, Baeken C, Benninger DH, Brunelin J, Di Lazzaro V, et al. Evidence-based guidelines on the therapeutic use of repetitive transcranial magnetic stimulation (rTMS): an update (2014–2018). Clinical Neurophysiology 2020;131:2150–206. 10.1016/j.clinph.2019.11.002.

[3] Ziemann U, Siebner HR. Inter-subject and intersession variability of plasticity induction by non-invasive brain stimulation: boon or bane? Brain Stimul 2015;8:662–3. 10.1016/j.brs.2015.01.409.

[4] Vida RG, Sághy E, Bella R, Kovács S, Erdősi D, Józwiak-Hagymásy J, et al. Efficacy of repetitive transcranial magnetic stimulation (rTMS) adjunctive therapy for major depressive disorder (MDD) after two antidepressant treatment failures: meta-analysis of randomized sham-controlled trials. BMC Psychiatry 2023;23:545. 10.1186/s12888-023-05033-y.

[5] Bi GQ, Poo MM. Synaptic modification by correlated activity: Hebb’s postulate revisited. Annu Rev Neurosci 2001;24:139–66. 10.1146/annurev.neuro.24.1.139.

[6] Dan Y, Poo MM. Spike timing-dependent plasticity of neural circuits. Neuron 2004;44:23–30. 10.1016/j.neuron.2004.09.007.

[7] Zrenner C, Desideri D, Belardinelli P, Ziemann U. Real-time EEG-defined excitability states determine efficacy of TMS-induced plasticity in human motor cortex. Brain Stimul 2018;11:374–89. 10.1016/j.brs.2017.11.016.

[8] Bergmann TO, Lieb A, Zrenner C, Ziemann U. Pulsed facilitation of corticospinal excitability by the sensorimotor μ-alpha rhythm. Journal of Neuroscience 2019;39:10034–43. 10.1523/JNEUROSCI.1730-19.2019.

[9] Humaidan D, Xu J, Kirchhoff M, Romani GL, Ilmoniemi RJ, Ziemann U. Towards real-time EEG–TMS modulation of brain state in a closed-loop approach. Clinical Neurophysiology 2024;158:212–7. 10.1016/j.clinph.2023.12.006.

[10] Stenroos M, Koponen LM. Real-time computation of the TMS-induced electric field in a realistic head model. Neuroimage 2019;203:116159. 10.1016/j.neuroimage.2019.116159.

[11] Pankka H, Lehtinen J, Ilmoniemi RJ, Roine T. Enhanced EEG forecasting: a probabilistic deep learning approach [preprint]. BioRxiv 2024:2024.01.16.575836. 10.1101/2024.01.16.575836.

[12] Lieb A, Zrenner B, Zrenner C, Kozák G, Martus P, Grefkes C, et al. Brain-oscillation-synchronized stimulation to enhance motor recovery in early subacute stroke: a randomized controlled double-blind three-arm parallel-group exploratory trial comparing personalized, non-personalized and sham repetitive transcranial magnetic stimulation (acronym: BOSS-STROKE). BMC Neurol 2023;23:204. 10.1186/s12883-023-03235-1.

[13] Zhu X, Song B, Ni Y, Ren Y, Li R. Software defined anything—from software-defined hardware to software defined anything. Business Trends in the Digital Era, 2016, p. 83–103. 10.1007/978-981-10-1079-8_5.

[14] Hassan U, Pillen S, Zrenner C, Bergmann TO. The Brain Electrophysiological recording C STimulation (BEST) toolbox. Brain Stimul 2022;15:109–15. 10.1016/j.brs.2021.11.017.

[15] Mikkili S, Panda AK, Prattipati J. Review of real-time simulator and the steps involved for implementation of a model from MATLAB/SIMULINK to real-time. Journal of the Institution of Engineers (India): Series B 2015;96:179–96. 10.1007/s40031-014-0128-6.

[16] Ciliberti D, Kloosterman F. Falcon: A highly flexible open-source software for closed-loop neuroscience. J Neural Eng 2017;14:045004. 10.1088/1741-2552/aa7526.

[17] Gansler JS, Lucyshyn W. Commercial-off-the-shelf (cots): doing it right (Technical Report No. UMD-AM-08-129). 2008.

[18] Raschka S, Patterson J, Nolet C. Machine learning in Python: main developments and technology trends in data science, machine learning, and artificial intelligence. Information (Switzerland) 2020;11:193. 10.3390/info11040193.

[19] Larrucea X, Santamaria I, Colomo-Palacios R, Ebert C. Microservices. IEEE Softw 2018;35:96–100.

[20] Abeni L, Balsini A, Cucinotta T. Container-based real-time scheduling in the Linux kernel. ACM SIGBED Review 2019;16:33–8. 10.1145/3373400.3373405.

[21] Li S, Zhang H, Jia Z, Li Z, Zhang C, Li J, et al. A dataflow-driven approach to identifying microservices from monolithic applications. Journal of Systems and Software 2019;157:110380. 10.1016/j.jss.2019.07.008.

[22] Majors C, Fong-Jones L, Miranda G. Observability engineering. O’Reilly Media, Inc.; 2022.

[23] Jindal A, Podolskiy V, Gerndt M. Performance modeling for cloud microservice applications. Proceedings of the ACM/SPEC International Conference on Performance Engineering (ICPE), 2019, p. 25–32. 10.1145/3297663.3310309.

[24] Macenski S, Foote T, Gerkey B, Lalancette C, Woodall W. Robot Operating System 2: design, architecture, and uses in the wild. Sci Robot 2022;7:eabm6074. 10.1126/scirobotics.abm6074.

[25] Kay J, Tsouroukdissian AR. Real-time control in ROS and ROS 2.0. Proceedings of the ROSCon 2015, Hamburg, Germany: 2015.

[26] Blass T, Hamann A, Lange R, Ziegenbein D, Brandenburg BB. Automatic latency management for ROS 2: Benefits, challenges, and open problems. Proceedings of the IEEE Real-Time and Embedded Technology and Applications Symposium, RTAS, 2021, p. 264–77. 10.1109/RTAS52030.2021.00029.

[27] Andrist B, Sehr V, Garney B. C++ high performance: master the art of optimizing the functioning of your C++ code. Packt Publishing Ltd; 2020.

[28] Merkel D. Docker: lightweight Linux containers for consistent development and deployment. Linux Journal 2014;2014:2.

[29] Jia X, Zhou Y, Hussain Y, Yang W. An empirical study on Python library dependency and conflict issues. Proceedings of the IEEE 24th International Conference on Software Quality, Reliability and Security (QRS), 2024, p. 504–15.

[30] Cinque M, Corte R Della, Pecchia A. Microservices monitoring with event logs and black box execution tracing. IEEE Trans Serv Comput, 2022, p. 294–307. 10.1109/TSC.2019.2940009.

[31] Jangla K. Docker Compose. In: Accelerating development velocity using Docker: Docker across microservices, Springer; 2018, p. 77–98.

[32] Tolaram N. systemd. In: Software development with Go: cloud-native programming using Golang with Linux and Docker, Springer; 2022, p. 325–45.

[33] Reghenzani F, Massari G, Fornaciari W. The real-time Linux kernel: a survey on PREEMPT_RT. ACM Comput Surv 2019;52:1–36. 10.1145/3297714.

[34] Madden MM. Challenges using Linux as a real-time operating system. AIAA Scitech 2019 Forum, 2019, p. 0502. 10.2514/6.2019-0502.

[35] Fedosejev A. React.js essentials. Packt Publishing Ltd; 2015.

[36] Mili H, Elkharraz A, Mcheick H. Understanding separation of concerns. IEE Proceedings - Software 2004:75–84.

[37] Watt ST, Achanta S, Abubakari H, Sagen E, Korkmaz Z, Ahmed H. Understanding and applying precision time protocol. Proceedings of the Saudi Arabia Smart Grid (SASG), 2016, p. 1–7. 10.1109/SASG.2015.7449285.

[38] Kopetz H, Ochsenreiter W. Clock synchronization in distributed real-time systems. IEEE Transactions on Computers 1987;36:933–40. 10.1109/TC.1987.5009516.

[39] Cardwell N, Savage S, Anderson T. Modeling TCP latency. IEEE INFOCOM 2000;3:1742–51. 10.1109/infcom.2000.832574.

[40] Shajii A, Numanagić I, Leighton AT, Greenyer H, Amarasinghe S, Berger B. A Python-based programming language for high-performance computational genomics. Nat Biotechnol 2021;39:1062–4. 10.1038/s41587-021-00985-6.

[41] Kundu B, Vassilev V, Lavrijsen W. Efficient and accurate automatic Python bindings with cppyy C Cling [preprint]. ArXiv Preprint ArXiv:230402712 2023.

[42] Harris CR, Millman KJ, van der Walt SJ, Gommers R, Virtanen P, Cournapeau D, et al. Array programming with NumPy. Nature 2020;585:357–62. 10.1038/s41586-020-2649-2.

[43] Virtanen P, Gommers R, Oliphant TE, Haberland M, Reddy T, Cournapeau D, et al. SciPy 1.0: fundamental algorithms for scientific computing in Python. Nat Methods 2020;17:261–72. 10.1038/s41592-019-0686-2.

[44] Okuta R, Unno Y, Nishino D, Hido S, Loomis C. Cupy: a NumPy-compatible library for NVIDIA GPU calculations. Proceedings of the Workshop on Machine Learning Systems (LearningSys) at the 31st Annual Conference on Neural Information Processing Systems (NIPS), 2017.

[45] Abadi M, Barham P, Chen J, Chen Z, Davis A, Dean J, et al. TensorFlow: a system for large-scale machine learning. Proceedings of the 12th USENIX Symposium on Operating Systems Design and Implementation (OSDI 16), 2016, p. 265–83.

[46] Li J, Li J. Dual circular buffer architecture for digital FIR/IIR filters. Proceedings of the Midwest Symposium on Circuits and Systems (MWSCAS), 2021, p. 400–3. 10.1109/MWSCAS47672.2021.9531678.

[47] Hughes CJ. Single-instruction multiple-data execution. Morgan & Claypool Publishers; 2015. 10.2200/S00647ED1V01Y201505CAC032.

[48] Ward L, Pauloski JG, Hayot-Sasson V, Chard R, Babuji Y, Sivaraman G, et al. Cloud services enable efficient AI-guided simulation workflows across heterogeneous resources. Proceedings of the IEEE International Parallel and Distributed Processing Symposium Workshops (IPDPSW), 2023, p. 32–41. 10.1109/IPDPSW59300.2023.00018.

[49] Prytz G, Johannessen S. Real-time performance measurements using UDP on Windows and Linux. Proceedings of the IEEE International Conference on Emerging Technologies and Factory Automation (ETFA), 2005. 10.1109/etfa.2005.1612771.

[50] Koponen LM, Nieminen JO, Ilmoniemi RJ. Multi-locus transcranial magnetic stimulation—theory and implementation. Brain Stimul 2018;11:849–55. 10.1016/j.brs.2018.03.014.

[51] Nieminen JO, Sinisalo H, Souza VH, Malmi M, Yuryev M, Tervo AE, et al. Multi-locus transcranial magnetic stimulation system for electronically targeted brain stimulation. Brain Stimul 2022;15:116–24. 10.1016/j.brs.2021.11.014.

[52] Kahilakoski O-P, Sinisalo H, Nieminen JO, Alkio K, Valén K, Kozák G, et al. A high-precision timing method and digital interface for closed-loop TMS. Submitted to biorXiv.

[53] Lyon BK, Popov G. Managing risk through layers of control. Prof Saf 2020;65:25–35.

[54] Rossi S, Antal A, Bestmann S, Bikson M, Brewer C, Brockmöller J, et al. Safety and recommendations for TMS use in healthy subjects and patient populations, with updates on training, ethical and regulatory issues: expert guidelines. Clinical Neurophysiology 2021;132:269–306. 10.1016/j.clinph.2020.10.003.

[55] Zrenner C, Galevska D, Nieminen JO, Baur D, Stefanou MI, Ziemann U. The shaky ground truth of real-time phase estimation. Neuroimage 2020;214:116761. 10.1016/j.neuroimage.2020.116761.

